# Single Particle Adsorption and Response Quantification for Functional AAVx Titer and Dose Analysis

**DOI:** 10.64898/2026.05.28.728583

**Authors:** Chiranth K. Nagaraj, Kim Truc Nguyen, Kwang Joo Kwak, Bilal Bayazit, Xin Huang, Jackson Newell, Jacob Doon-Ralls, Xilal Y. Rima, Scott Q. Harper, Nizar Y. Saad, Eduardo Reátegui

**Affiliations:** William G. Lowrie Department of Chemical and Biomolecular Engineering, The Ohio State University, Columbus, OH, 43210, USA; Spot Biosystems, Columbus, OH, 43210, USA; Jerry R. Mendell Center for Gene Therapy, Abigail Wexner Research Institute at Nationwide Children’s Hospital, Columbus, OH, 43215, USA; Diabetes and Metabolism Research Center, Division of Endocrinology, Diabetes, and Metabolism, Department of Internal Medicine, The Ohio State University Wexner Medical Center, Columbus, OH, 43210, USA; Department of Pediatrics, The Ohio State University College of Medicine, Columbus, OH, 43210, USA; Comprehensive Cancer Center, The Ohio State University; Columbus, OH 43210, USA

**Keywords:** Single-particle, AAV, gene therapy, diffraction-limited imaging, Simulation

## Abstract

Accurate quantification of functional adeno-associated virus (AAV) dose remains a critical limitation in gene therapy, where titers are defined using ensemble measurements of viral genomes or capsid concentration that average across heterogeneous populations and weakly predict transduction. We present single-particle adsorption and response quantification (SPARQ), a diffraction-limited imaging platform for quantifying AAVx vectors, regardless of serotype, at single-nanoparticle and single-cell resolution. SPARQ immobilizes and resolves individual AAVx, enabling concurrent measurement of capsid-associated and genome-associated signals before cell exposure. Across multiple serotypes, including AAV2, AAV5, and AAVrh74, SPARQ identifies full-capsid fractions, consistent with bulk measurements obtained by ddPCR/ELISA, DLS/UV-Vis, charge detection, and mass photometry, while reporting narrower distributions than ensemble-derived values and revealing systematic discrepancies in bulk-derived titers arising from population averaging. Surface immobilization is achieved by charge-dependent AAVx-surface interactions, as supported by multiphysics simulations, which capture serotype-dependent adsorption kinetics governed by capsid electrostatics for AAV2, AAV5, AAV6, and AAV9. Furthermore, SPARQ directly measures transduction as a function of particle number and reveals a threshold response. Transduction efficiency increased from ∼29% (S.D ± 3.31%) at ∼18 (S.D ± 7.55) AAVs per cell to ∼66% (S.D ± 1.66%) at ∼42 (S.D ± 9.69) AAVs per cell, which is comparable with conventional in vitro assay requiring 10^5^ AAVs/cell. Cumulative fluorescence measurements across large micro-patterned arrays of single cells recapitulate bulk-like high-throughput scaling, while SPARQ’s single-nanoparticle resolution reveals functional heterogeneity that is masked by population averaged assays. SPARQ provides a particle-resolved framework for functional AAVx titration and quantitative characterization, enabling the development of quality control and dosing strategies in gene therapy manufacturing.

**Teaser:** SPARQ combines light-activated surface viral adsorption with diffraction-limited imaging to directly quantify AAV integrity, heterogeneity, and functional transduction at the single-particle level.

## Introduction

Adeno-associated viruses (AAVs) have emerged as leading vectors for in vivo gene delivery, enabling durable transgene expression across a growing range of therapeutic indications. Clinical success across neuromuscular, retinal, and central nervous system disorders has accelerated the translation of AAV-based gene therapies^1,2,3,4^, while simultaneously exposing fundamental limitations in how viral dosing, vector quality, and functional AAV availability are currently defined in the field. In both preclinical and clinical contexts, AAV dosing is predominantly specified using ensemble measurements, such as viral genome copy number, capsid concentration, or nominal multiplicity of infection (MOI) that average across large and heterogeneous viral populations^5,6,7,8^. These bulk metrics implicitly assume that all viral particles contribute equivalently to transduction, despite mounting evidence that AAV preparations exhibit substantial particle-to-particle variability in capsid composition, genome packaging, structural integrity, and biological function^4,7,9,10^.

AAV transduction is a multistep and intrinsically probabilistic process, encompassing capsid binding, cellular uptake, endosomal escape, nuclear trafficking, uncoating, and transcriptional activation^11,12,13^. Failure at any of these stages makes an individual AAV functionally inert, even if it is counted as part of the delivered dose. Prior studies have shown heterogeneity in empty versus genome-containing capsids, serotype-dependent differences in capsid stability and transient capsid disassembly kinetics, stochastic genome release dynamics, and variability in intracellular trafficking efficiency^9,14–18^. Self-complementary AAVs partially mitigate second-strand synthesis limitations but do not eliminate upstream stochastic barriers^19^. Yet, existing analytical workflows, including qPCR, ddPCR, SEC-MALS, ELISA-based capsid quantification, analytical ultracentrifugation, and bulk infectivity assays^20–26^ are fundamentally limited to population-averaged outcomes. As a consequence, bulk titers often correlate poorly with the probability of how many viral particles are physically available to a given cell, whether those particles retain intact genomes at the time of delivery, or the probability of how particle-level variability translates into functional gene expression^27^.

Recent advances in single-particle characterization have begun to reveal the extent of nanoscale heterogeneity within viral and extracellular vesicle populations^9,28–31^. High-throughput optical and biophysical methods have enabled direct measurement of particle size, diffusivity, binding affinity, and cargo content at the individual virion level, challenging long-standing assumptions derived from ensemble measurements. However, these approaches remain largely descriptive: while they provide great insight into particle composition, they do not establish links between particle number, particle integrity, and biological outcome. In particular, a fundamental question remains unresolved: the probability of how many AAV particles are functionally required to produce gene expression in a single cell? Addressing this question experimentally requires the ability to both quantize viral particle numbers and observe cellular responses under controlled single-cell conditions.

Here, we introduce Single-Particle Adsorption and Response Quantification (SPARQ), a diffraction-limited imaging platform for quantifying the functional titer of AAVx vectors. Using total internal reflection fluorescence microscopy (TIRFM), SPARQ resolves individual AAVx particles and concurrently quantifies capsid and genome-associated fluorescence signals, enabling single-AAV level identification of genome-containing, genome-compromised, and empty viral vectors before cellular exposure. In addition, SPARQ uses UV-light patterning to enable electrostatic adsorption of AAVx, allowing individual AAV particles to be immobilized on predefined surface patterns with controlled spatial geometry. We further show that AAV adsorption exhibits serotype-dependent binding dynamics governed by capsid electrostatics, with experimentally measured kinetics^32^ quantitatively captured by multi-physics simulations. By decoupling viral adsorption from bulk suspension delivery, SPARQ enables direct counting of AAVx particles at pre-defined locations and systematic analysis of nanoparticle-surface interactions. Single-cell patterning reveals that effective transduction is threshold-governed. Unlike ensemble assays, this platform enables assigning a defined number of viral particles to a single cell and directly measuring the resulting probability of transduction. SPARQ provides a functional definition of titer grounded in particle-resolved imaging and biological outcome, providing a generalizable strategy for accurate viral dosing, quality assessment, and fundamentals of gene delivery processes.

## Materials and Methods

### SPARQ platform fabrication and viral adsorption patterning

SPARQ devices were fabricated using gold-coated coverslips for AAVx titer quantification, and UV micropatterned coverslips for single-cell AAVx transduction assay. Briefly, high-precision borosilicate glass coverslips (24 × 75 × 0.15 mm, D 263® M; Schott AG, Mainz, Germany) were thoroughly cleaned in an ultrasonic bath, first in ethanol and then in deionized (DI) water, for 5 min each. This cleaning process was repeated, after which the coverslips were dried with nitrogen gas. Next, the coverslips were treated in a UV-ozone cleaner (Jelight, Irvine, CA) for 15 mins. A 2 nm titanium film was deposited on the cleaned coverslips using electron beam evaporation (DV-502A; Denton Vacuum, Moorestown, NJ) to promote adhesion of a subsequent gold layer. Using the same technique, a 12 nm gold layer was applied over the titanium. The resulting gold-coated coverslips were dried with nitrogen gas and secured to a 64-well ProPlate® microarray system (Sigma-Aldrich, St. Louis, MO). For AAV adsorption experiments, 20 µL of AAVx samples were added to each well and incubated for 45 mins before imaging. For controlled immobilization of viral particles, SPARQ substrates were further prepared by adapting the light-induced extracellular vesicles and nanoparticles adsorption (LEVA) workflow reported previously^33^. Briefly, glass coverslips were washed thrice with ethanol/DI-water and sonicated, then they were plasma-treated for 1 min, followed by incubation with 0.01% (w/v) poly-L-lysine (PLL; 2h), rinsing in 0.1 M HEPES, and passivation with methoxy-PEG-SVA (mPEG-SVA; 100 mgmL⁻¹ in 0.1 M HEPES, pH 8.5; 2h) before DI-water rinsing and drying under nitrogen. A PLPP photo-activator layer was subsequently applied, and patterns were etched using a digital micromirror device (DMD) microscope module (PRIMO, Alvéole) on an inverted microscope (Nikon Eclipse Ti2), enabling maskless UV exposure of 30 mJmm⁻² to locally photoetch the PEG layer and generate positively adsorptive regions. The pattern template consisted of 15 µm-diameter circular features arranged at 50 µm center-to-center spacing in a 10 × 10 × 15 array, which was uploaded to the Leonardo version 5 PRIMO software and projected onto the functionalized surface to generate the SPARQ patterned microarray.

### Immunofluorescence labeling of AAVx capsids and genomes

A 3% (w/v) BSA solution in PBS was incubated in each well at room temperature for 1 h to block nonspecific interactions. After withdrawing the BSA solution, 1 µgmL^-1^ solution of the fluorescently conjugated antibodies in 10 % (w/v) normal goat serum (NGS; Thermo-Fisher Scientific, Waltham, MA) was incubated in the wells for 1 h at room temperature in a dark environment **(Table S2)**. Excess antibodies were rinsed and removed by passing through a size exclusion chromatography column (qEV Single 35 nm IZON Inc, Boston, MA) and by incubating in PBS for 5 mins, then pipetting up and down the solution 10×. The rinsing step was repeated 3× in a dark environment. For genome staining, 100 µL of purified AAV samples at 1×10^11^ vgmL^-1^ were diluted in 100 × buffer to stabilize nucleic acids in capsids. Ethidium homodimer-1 (EthD-1) (Thermo-Fisher Scientific, Waltham, MA) was added to a final working concentration of 5 µM. Following incubation, excess EthD-1 was washed by following the previously mentioned steps under the Immunofluorescence labeling of AAVx capsid section. After washing, the 20 µL of AAVx samples were immediately added to SPARQ for diffraction-limited imaging.

### Diffraction-limited image acquisition and processing

A 10×10 array of images was acquired via TIRFM (Nikon, Melville, NY) at 596×596 pixels (px) for analyzing single AAVx signals and at 1192×1192 px for larger region of interest (ROI) acquisitions for each well with a 100× oil immersion objective. An exposure time of 200 ms and a laser power of 40% were maintained across all channels to ensure assay consistency. TIRFM images were quantified by measuring the total and mean fluorescence intensities of each TIRFM-detected spot. Histograms were generated from the total and mean intensities of individual spots, and scatter plots were generated from the mean intensity and size of single spots detected by TIRFM. Relative and total fluorescence intensities of the sample were obtained from custom-built algorithms that were previously reported in the literature ^30,31^.

### Transmission electron microscopy (TEM) for AAVx particles

Two 20 µL droplets of water for injection (WFI) and two 20 µL droplets of negative stain (uranyLess EM stain, Electron Microscopy Sciences, Hatfield, PA) were placed on a strip of parafilm. The TEM grid was subjected to plasma treatment for 1 min. Then, 10 µL of AAVx solution was carefully placed onto the treated surface of the grid. The virion solution was incubated on the surface of the grid for 1 min and gently blotted with filter paper to remove excess liquid. These grids were submerged in the WFI droplet and blotted dry using filter paper. The process was repeated with the second WFI droplet. The grids were submerged in the first droplet of the negative stain, followed by blotting, and then submerged again in the second droplet of the negative stain. The grid was allowed to incubate in the stain for approximately 22 secs before gently wicking away the excess solution using filter paper. To ensure thorough drying, the stained grids were stored in a grid box overnight. Afterward, TEM imaging was performed using a Tecnai TF-20 microscope (FEI Company, Hillsboro, OR) operating at 200 kV.

### AAVx vector genome titration by qPCR and ddPCR with DNase treatment

Single-stranded (ss) and self-complementary (sc) AAV9.CMV.eGFP.U6.miLacZ, ssAAV6.CMV.eGFP, and ss and scAAVrh74.CMV.eGFP.U6.milacZ with enhanced green fluorescent protein (eGFP) reporter was titered by quantitative PCR using a plasmid standard curve and a CMV primer/probe set (CMV forward primer: TGGAAATCCCCGTGAGTCAA, CMV reverse primer: CATGGTGATGCGGTTTTGG, and TaqMan probe sequence: CCGCTATCCACGCCCATTGATG). Briefly, serial viral sample dilutions at a 1:10 dilution factor were prepared. The number of serial dilutions depends on the virus concentration and can range between 2 serial dilutions at 1:10^4^ and 1:10^5^ and 4 serial dilutions (1:10^2^, 1:10^3^, 1:10^4^ and 1:10^5^). Before running the qPCR on the Bio-Rad CFX Connect, Real-Time system, diluted samples were treated with DNase I (Invitrogen Cat# 18047-019) at for 30 mins at 37° C, followed by 10 mins at 90° C, then a hold at 4° C. Samples were then treated with 1 µL of Proteinase K (Qiagen Cat# 19131 (stock conc.: 20 mgmL^-1^)) for 60 mins at 50° C, followed by 20 mins at 95° C and a hold at 4° C. Sample titers were calculated and reported as DNAse Resistant Particles (DRP). All used viruses were re-titered using the AAV ITR-2 probe (Bio-Rad, Expert design assay unique ID: dEXD15274642) on the Bio-Rad droplet digital PCR (ddPCR) and following Bio-Rad’s AAV vector genome ddPCR titration protocol. The ddPCR titers were similar to the CMV probe-based titration. To investigate the presence of capsid-less genome in the AAV preparations, AAV samples were DNase-treated without lysis before ddPCR titration was performed. Briefly, AAV samples were treated with Ambion DNase I (Invitrogen, Cat# AM2222) at 200 UmL^-1^ for 30 min at 37° C. Untreated samples were diluted similarly with a poly A+ solution (10 mM Tris-HCl, pH 8, 0.1 mM EDTA, 100 μgmL^-1^ poly[A], 0.01% Pluronic F-68 (Gibco, Cat# 24040032)). Samples were then serially diluted down an eight-tube strip (2 μl in 18 μl) using the poly A+ solution. The diluted viruses were lysed at 95° C for 10 mins using a standard thermal cycler. From each dilution, 5 μL was added to a mixture of ddPCR supermix for probes (no dUTPs) (Bio-Rad, Cat# 1863023) and AAV ITR-2 probe, each at 1× concentration. For droplet generation, 20 μL of the 25 μL reaction was transferred to a 96-well plate, heat-sealed with foil, and loaded into the QXDx automated droplet generator (Bio-Rad). The new droplet-containing 96-well plate was sealed, and amplification was performed using a C1000 touch thermal cycler (Bio-Rad) with the following conditions: 10 min at 95° C followed by 40 cycles of 30 seconds at 94° C and 1 min at 60° C followed by 10 min at 98° C. The PCR plate was subsequently read on a QX200 droplet reader (Bio-Rad). The data was processed using QX Manager software (Bio-Rad). Concentration values (copies μL^-1^) for each dilution were converted to stock vgμL^-1^ by multiplying each value by its respective total dilution factor (dilutions for DNase treatment, PCR reaction, and each serial dilution). The presented values are the averages of three dilutions that yield linear concentration values.

### AAVx capsid analysis by ELISA

ss, sc, and empty AAV9 capsids (kit control) were quantified by using the AAV9 titration ELISA kit (Progen, #PRAAV9). All samples were diluted to concentrations of 1×10^11^, 5×10^10^, 1×10^10^, 1×10^9^, 5×10^8^, 2.5×10^8^, 1.25×10^8^, and 6.13×10^7^ vgmL^-1^ with 1× assay buffer. A total of 100 µL of each sample was added to the ELISA plate and incubated at 37° C for 1 h. The wells were then washed three times in 1× assay buffer and incubated with 100 µL 1× biotin at 37 °C for 1h. After repeating the washing step, 100 µL 1× strep-HRP was added to the wells, and the plate was incubated at 37° C for 1 h. After another washing cycle, 100 μL of tetramethylbenzidine was added to the wells, and the plate was incubated at room temperature for 15 mins. The color reaction was stopped by adding 100 μL of stop solution, and the color intensity was measured at a wavelength of 450 nm using a photometer (SpectraMax iD5, Molecular Devices).

### AAVx adsorption kinetic simulations

COMSOL Multiphysics (v6.1, COMSOL Inc., Burlington, MA) licensed through the Ohio Supercomputer Centre (OSC, Columbus, OH) was used to model the transport and adsorption dynamics of nanoscale viral particles using a pre-defined light-induced extracellular vesicle and particle adsorption model^33^, by using 15 µm circle pattern with 50 µm apart center-to-center, and by using AAV6, AAV9, rAAV5, and AAV2 with zeta potentials -10 mV (S.D±0.45 mV)^34^, -20 (S.D±1.65 mV)^35^, -17.7 mV (S.D±1.9 mV)^36^, -3.5 mV (S.D±0.55 mV)^34^ respectively. The second simulation of full and empty AAVs on the gold SPARQ was performed as follows: the gold-coated SPARQ (3.5×3.5 mm) surface was defined as the binding substrate. AAV particles were modelled as spherical entities characterized by full and empty rAAV5 samples, with a zeta potential of −7.7 mV (S.D±2.2 mV) for the empty rAAV5, and a hydrodynamic diameter of ∼25 nm^36^. The electrostatic potential near the gold surface was solved using the Poisson-Boltzmann equation, accounting for the charge distribution induced by the gold and the surrounding ionic environment with PBS solution. The gold surface was modelled with a surface charge density influenced by the applied potential and ionic strength of the medium. Fluid flow within the micro-well was simulated using the Navier-Stokes equations under laminar flow conditions, while AAV particle motion was governed by the Langevin equation and particle tracing module, incorporating contributions from Brownian motion, hydrodynamic drag, and electrostatic interactions. Boundary conditions included a defined surface potential for the gold substrate and a no-slip condition for the fluid flow along the walls. The initial condition was specified as a uniform particle concentration of 2×10^9^, 1×10^9^, and 2×10^8^ AAVs mL^-1^, while the walls maintained constant pressure to ensure continuity. Mesh refinement was conducted near the gold surface to resolve the steep gradients in particle concentration and binding kinetics accurately. Time-dependent simulations were performed to track particle trajectories and surface binding dynamics. The adsorption profiles were analyzed to evaluate the effects of variables such as AAV kinetic energy, surface charge density, and ionic strength on binding efficiency. Simulation results were validated against experimental data obtained using fluorescence-tagged AAVs interacting with patterned surfaces and gold surfaces in controlled SPARQ and TIRFM imaging setups.

### Refeyn mass photometry

AAVx samples were diluted 10× in sterile-filtered PBS (pH 7.4) and equilibrated to room temperature before measurement. Mass photometry was performed using a SamuxMP mass photometer (Refeyn Ltd., Waltham, MA). Measurements were conducted according to manufacturer-recommended AAV workflows, targeting a final particle concentration of 1×10^11^ particles mL^-1^. For each sample, movies of individual particle landing events were recorded using AcquireMP software for 60 secs. Movies were analyzed in DiscoverMP v2024R2.1 to convert interferometric contrast into calibrated mass distributions. AAV populations were segmented using the built-in AAV overlay into empty, partial, and full regions, with region boundaries positioned to match the corresponding peaks in each calibrated histogram.

### HEK293T cells transduction

HEK293T cells with the passage number <10 was transduced with patterned AAV6 and maintained in Dulbecco’s Modified Eagle Medium (DMEM, Gibco, Grand Island, NY) supplemented with 10% (v/v) fetal bovine serum (FBS, Gibco, Grand Island, NY) and 1% penicillin-streptomycin (PS, Gibco, Grand Island, NY).

### Statistical Analysis

Statistical analyses were performed using the JMP Pro 14 software (JMP, Cary, NC) and Excel (Microsoft, Redmond, WA), where statistical significances were determined with *p* < 0.05.

## Results

### SPARQ for AAVx titer quantification

SPARQ is a diffraction-limited in situ single-particle imaging platform designed to quantify functional AAV titer by resolving individual AAV particles and their associated genomic content. To establish this particle-resolved framework, AAVs were dual-labeled to report capsid (VP1-3) and genome-associated signals for TIRFM imaging **(Fig. 1a-c)**. Individual fluorescent signals corresponding to single AAVx were captured within a selected ROI, with co-localized capsid and genome signals confirming intact AAVs **(Fig. 1c, d)**. Single-AAV fluorescence intensity distributions exhibited discrete, right-skewed profiles characteristic of individual nanoscale emitters rather than ensemble aggregates **(Fig. 1e, f)**. Quantification of capsid-genome co-localization demonstrated that the majority of AAVs retained genome-associated signals, confirming that SPARQ preserves viral integrity during surface capture **(Fig. 1g)**. Diffraction-limited fluorescence imaging shows a monotonic, titer-dependent increase in surface-bound AAV signal from 1×10^9^ to 1×10^11^ a.u. with negligible background in negative controls, and single-particle intensity distributions are clearly separated from PBS, confirming specific detection. TEM and cryo-EM further validated particle morphology and size, consistent with intact AAV capsids **(Fig. 1h).** These data establish a robust optical definition showing individual AAVs and their internal content.

**Figure 1:**
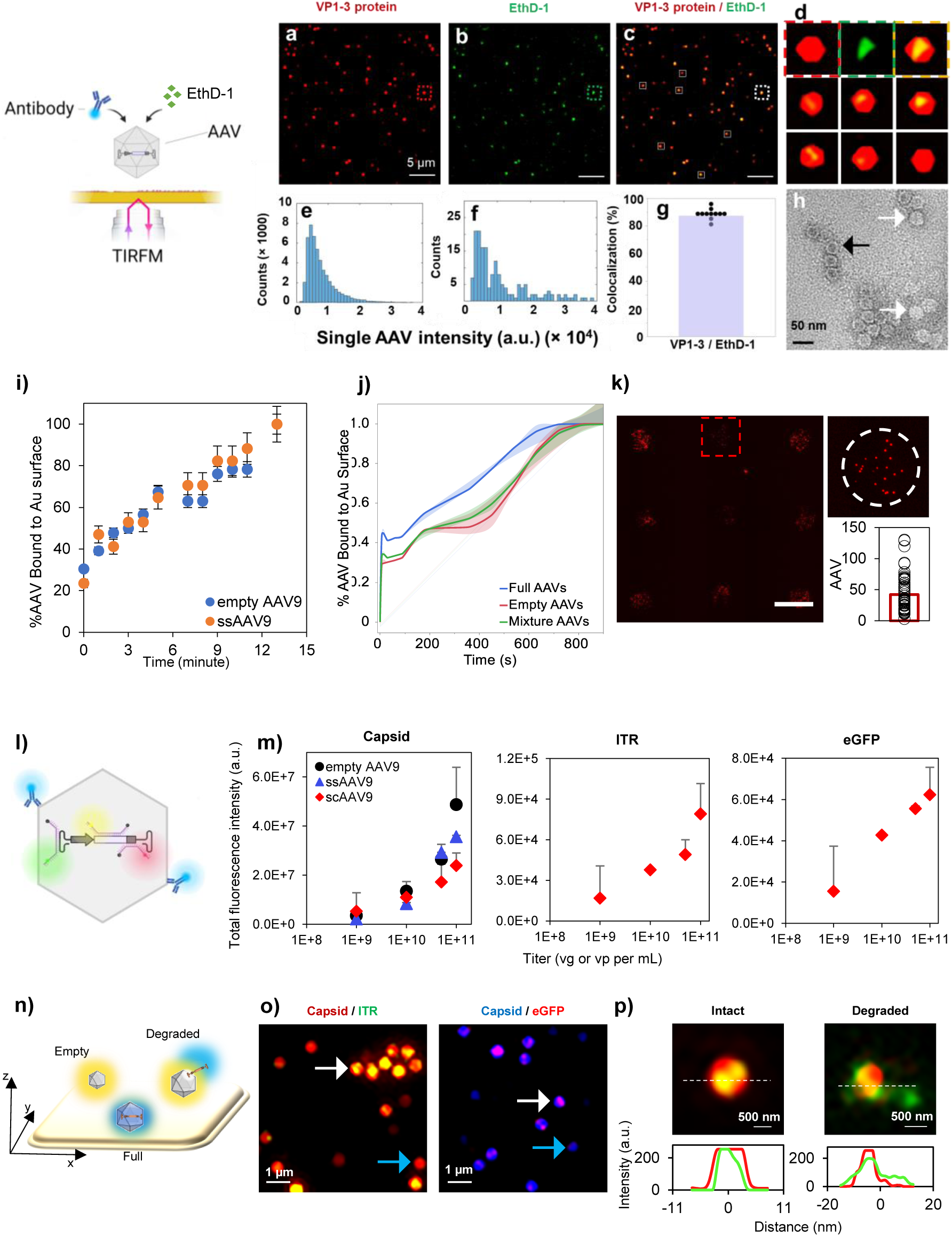
Simultaneous detection of DNA and protein capsid on a single AAVx particle with surface binding dynamics. Schematic of the experimental setup illustrating total internal reflection fluorescence microscopy (TIRFM) based detection of individual AAV particles using a fluorescent antibody. **(a)** Representative TIRFM images of VP1-3 capsid protein signal (red), **(b)** EthD-1 nucleic acid signal (green), and **(c)** merged channel co-localization images **(d)** demonstrating co-localization of capsid and genome signals with zoomed regions highlighting full (co-localized), empty (capsid-only), and degraded particles. Scale bars, 5 µm (main panels, n = 9). Histogram representation of single-particle fluorescence intensities for **(e)** capsid and **(f)** genome signals. **(g)** Quantification of capsid–genome co-localization efficiency using 100 images from distinct locations. Bars represent the mean 87 % (S.D ± 6.9 %; n = 3). Statistical significance assessed by two-sided unpaired Student’s *t*-test (*p* < 0.05). **(h)** Representative transmission electron microscopy (TEM) image confirming intact AAV morphology and particle integrity. White arrows indicate full AAVs; black arrows indicate empty AAVs. Scale bar, 50 nm. **(i)** Time-dependent experimental binding kinetics of AAVrh74 particles using SPARQ, showing increased surface occupancy over time (n = 3). **(j)** Normalized cumulative binding curves comparing full AAVs, empty AAVs, and mixture populations from simulation (n = 3). **(k)** Representative TIRFM field of view illustrating the spatial distribution of surface-bound AAVs. Right, quantification of particle counts per field with mean - 42.27 (S.D ± 9.69; n = 36; one-way ANOVA with Tukey’s HSD test)**. (l)** Schematic illustrating classification of AAV particles based on capsid and genome fluorescence signatures into full, empty, and degraded populations. **(m)** Total fluorescence intensity analysis of capsid signal as a function of viral titer for empty, full, and standard AAV preparations (n = 3). **(n)** Schematic depicting differential surface behavior of empty, full, and degraded AAVs under TIRFM illumination. **(o)** Dual-color TIRFM images showing capsid (red) and genome (blue) channels for intact particles (white arrows) and partially degraded particles (cyan arrows). Scale bars, 1 µm (n = 3). **(p)** Representative line-scan intensity profiles across intact (left) and degraded (right) particles, demonstrating reduced genome signal in degraded capsids (n = 10).

### SPARQ enables time-resolved quantification of AAV adsorption kinetics

AAVx adsorption kinetics were evaluated by time-dependent continuous detection of AAVx capsid protein and genome co-localization. Time-lapse TIRFM imaging revealed a monotonic increase in surface-bound AAVs following exposure to viral suspensions, with adsorption progressing rapidly during the initial minutes and approaching saturation thereafter **(Fig. 1i)**. Importantly, multi-physics adsorption kinetics differed between empty and genome-containing AAVs, with full AAVs exhibiting higher binding efficiencies and faster accumulation on the SPARQ surface compared to empty capsids **(Fig. 1j)**. The simulated SPARQ surface showed that the occupancy increased in a concentration-dependent manner, allowing direct quantification of the number of AAVs bound per unit surface area over fixed endpoint time and the trend matched with the experimental outcomes. The maximum surface concentration observed was 3.2×10^10^ particles/micro-well, corresponding to the theoretical monolayer coverage limit without particle stacking. For empty rAAV5s (charge: −7.7 mV, S.D±2.2 mV), binding exhibited an exponential increase in surface coverage, driven predominantly by electrostatic interactions. The binding saturated at 12 mins and 43 secs. Statistical analysis revealed consistency across simulations, with an average binding rate of 0.42 (S.D±0.04) particles (1×10^5^)/m^2^/s during the exponential phase. Statistical analysis showed consistency across replicates, with a coefficient of variation (CV) of 8.9% (*p* < 0.01, one-way ANOVA). For full AAVs (charge: −17.7 mV, S.D±1.9 mV), saturation was achieved within 10 mins and 28 secs, approximately 18 % faster than empty AAVs (*p* < 0.01, paired t-test). The binding rate during the exponential phase was significantly higher at 0.56 (S.D ± 0.05) particles (1×10^5^)/m^2^/s with a CV of 7.2%. In the 50:50 mixture of full and empty AAVs, the binding profile displayed a two-phase trend. Full AAVs bound initially at a faster rate, dominating the early exponential phase, followed by the gradual binding of empty AAVs. The maximum surface coverage remained 3.2×10^10^ particles/micro-well, with saturation occurring in 13 mins and 33 secs, approximately 2.5 mins longer than empty AAVs alone. Particle-particle interactions and overlapping influenced the delayed saturation. The mixed binding rate was split into two phases: 0.51 (S.D±0.04) particles/m^2^/s for full AAVs and 0.37 (S.D±0.03) particles/m^2^/s for empty AAVs during their respective dominant binding periods (*p* < 0.05, mixed-model analysis). The overall CV for the mixture simulations was 6.5% despite the complexity of competitive binding. No significant binding was observed in control simulations using PBS with no particles or with cationic particles (∼25 nm) on the gold surface. For coverslips (non-gold substrates), binding was minimal (< 10% surface coverage at 5 mins) and did not exhibit exponential trends. This showed the influence of the SPARQ surface in mediating AAV binding through electrostatic interactions.

Furthermore, at a steady state, SPARQ displayed discrete clusters of immobilized AAV9s, enabling direct counting of the AAV number per pattern. Quantitative analysis revealed an average occupancy of ∼42 (S.D ± 9.69 AAV9s) per pattern under standard experimental conditions. Notably, control regions outside patterned areas exhibited negligible particle accumulation, confirming that adsorption is spatially confined and governed by surface patterning rather than nonspecific deposition **(Fig. 1k)**. Thus, SPARQ is a quantitative platform for generating well-defined viral microarrays with known AAVx numbers per addressable location. Also, these measurements demonstrate that SPARQ provided deterministic control over AAV immobilization while preserving the stochastic nature of individual adsorption events.

### SPARQ decouples capsid abundance, genome availability, and functional output

To examine how bulk viral input translates into a single AAV level signal, we quantified capsid-associated fluorescence, inverted terminal repeat (ITR) targeted genome-associated fluorescence signal, and reporter expression across increasing AAV titers using diffraction-limited single-particle imaging **(Fig. 1l)**. To assess variation in viral genome configuration and functional titer, we performed diffraction-limited imaging analysis across empty AAV9, single-stranded AAV9 (ssAAV9), and self-complementary AAV9 (scAAV9) preparations over a range of input concentrations spanning 10⁹-10¹¹ vgmL⁻¹. Across all titers, empty AAV9 preparations exhibited the highest measured signal intensity, increasing from ∼3.5×10⁶ at 10⁹ vgmL⁻¹ to ∼4.9×10⁷ at 10¹¹ vgmL⁻¹, reflecting accumulation of capsid-associated signal in the absence of packaged genomes. In contrast, ssAAV9 displayed consistently lower signal intensities at matched inputs, rising from ∼2.1×10⁶ to ∼3.6×10⁷ over the same titer range, indicating a compressed dynamic range relative to empty capsids. Self-complementary AAV9 (scAAV9) exhibited an intermediate but systematically lower response, reaching ∼2.4×10⁷ at 10¹¹ vgmL⁻¹, with reduced variance across all titers **(Fig. 1m)**. Furthermore, the capsid-associated fluorescence increased monotonically with input concentration, showing the steepest scaling among the three samples and reflecting accumulation of surface-immobilized AAVs. In contrast, the ITR signal increased more modestly with titer, exhibiting a reduced slope relative to the capsid signal and indicating that increases in particle number do not translate proportionally into genome-competent particles. Notably, enhanced green fluorescent protein (eGFP) expression exhibited scaling behavior that followed the ITR signal rather than capsid abundance. While the eGFP signal increased with input concentration, its dynamic range was substantially compressed relative to capsid measurements, indicating that functional output was constrained by genome availability rather than by total AAV count. Furthermore, the total fluorescence intensity of AAVs decreases with increasing temperature, indicating temperature-dependent loss of capsid-associated signal consistent with reduced structural stability at elevated temperatures. In contrast, bulk absorbance measurements for empty AAV9, ssAAV9, and scAAV9 scale monotonically with capsid concentration across the same range. Therefore, highlighting increasing decoupling between nominal viral input and functional outcome at higher concentrations, which further indicates substantial AAV-to-AAV heterogeneity.

### SPARQ enables colocalization of protein and DNA signals in single AAVx for determining capsid integrity

AAV co-localization analysis revealed the presence of empty capsids, genome-containing particles, and genome-degraded or partially compromised particles within the same preparation **(Fig. 1n)**. Diffraction-limited imaging of individual AAVs enabled direct assessment of capsid presence relative to genome-associated signal **(Fig. 1o)**. Two-dimensional scatter plots of single particle mean intensity versus effective particle area reveal distinct clustering behaviors for empty AAV, ssAAV, and scAAV across capsid (3D), eGFP, and ITR channels. Dual-fluorescent diffraction-limited imaging shows co-localization of VP1-3 capsid signal with ITR and eGFP signals for genome-containing AAV particles, whereas empty AAV preparations exhibit capsid signal in the absence of detectable genome-associated fluorescence. Co-localization of capsid and ITR fluorescence identified intact viral vectors, whereas particles displaying capsid signals with spatially offset genome signals were classified as genome compromised. Within the same preparation, both intact and compromised particles were observed, demonstrating substantial heterogeneity in genome retention at the single-particle level. Capsid-only AAVs, indicative of empty capsids, were also easily observed and distinguished from genome-containing viral vectors based on the absence of ITR or reporter-associated signal. Total fluorescence intensity measured for capsid (3D), eGFP, and ITR fluorescent channels scales differently with increasing viral input, with capsid signal exhibiting the steepest and most linear increase, while ITR targeted and reporter (eGFP) signals display compressed and non-proportional scaling. Line-scan analysis across individual particles further resolved structural differences between intact and degraded viral vectors **(Fig. 1p)**. Diffraction-limited imaging across multiple axial planes (−40 to +80 nm) enabled reliable detection of genome-associated fluorescence within individual AAVx particles despite axial step sizes exceeding the physical capsid diameter. This is enabled by the extended axial point-spread function, which integrates fluorescence over a depth larger than the viral vectors, allowing genome detection independent of particle orientation on the surface. Apparent particle sizes exceed the true capsid dimensions due to convolution with the optical point-spread and reflect fluorescence distribution rather than physical size. Intact particles exhibited overlapping capsid and genome intensity profiles with coincident maxima and symmetric distributions, consistent with a retained genome within the capsid. In contrast, degraded AAVs displayed broadened and asymmetric genome-associated profiles that were spatially displaced relative to the capsid signal, indicating partial genome loss. These data demonstrated that diffraction-limited imaging resolves functional heterogeneity at the single-AAV level.

### Serotype-dependent binding dynamics and single-cell functional outcomes resolved by SPARQ

To examine how capsid properties influence surface binding, particle transport, and functional transduction, we compared multiple AAV serotypes using SPARQ for single AAV imaging, and single-cell transduction assays. Bulk fluorescence measurements revealed a monotonic increase in total AAV-associated signal with an increasing input titer for AAV2, AAV5, and AAVrh74, but with clear serotype-specific differences in absolute intensity at matched concentrations **(Fig. 2a)**. Imaging confirmed that these differences arose from distinct AAV distributions and capsid–genome co-localization patterns across serotypes **(Fig. 2b)**, rather than uniform increases in AAV number alone. To benchmark single-AAV measurements against commonly used ensemble assays, we compared the fraction of genome-containing (full) capsids across AAV2, AAV5, and AAVrh74 using SPARQ-AAV analysis, ddPCR/ELISA-based infectivity readouts, DLS/UV-vis, and charge-detection mass spectrometry (CDMS) **(Fig. 2c)**. SPARQ measurements yielded moderate and tightly distributed estimates of full capsid fraction, with mean values of 35.3% (S.D±2.3%) for AAV2, 38.5% (S.D±1.7%) for AAV5, and 44.9% (S.D±3.7%) for AAVrh74. In contrast, ensemble-based methods reported substantially higher but more variable values, with ddPCR/ELISA-derived estimates ranging from 20.8% (AAV2) to 71.4% (AAV5), DLS/UV-VIS reporting 58-84%, and CDMS reporting 75-85% across serotypes. Furthermore, mass spectrometry data showed that PBS controls show no detectable particulate signal, while mass photometry imaging showed discrete AAV particle populations with well-defined mass signatures and quantified distinct empty, partially filled, and full capsid populations and revealed differences in empty/full ratios between AAVrh74 preparations. Notably, the relative ordering of serotypes differed across methods. While SPARQ measurements showed a gradual increase in full capsid fraction from AAV2 to AAVrh74, bulk assays amplified inter-serotype differences and consistently overestimated full AAV abundance relative to TIRFM imaging.

**Figure 2:**
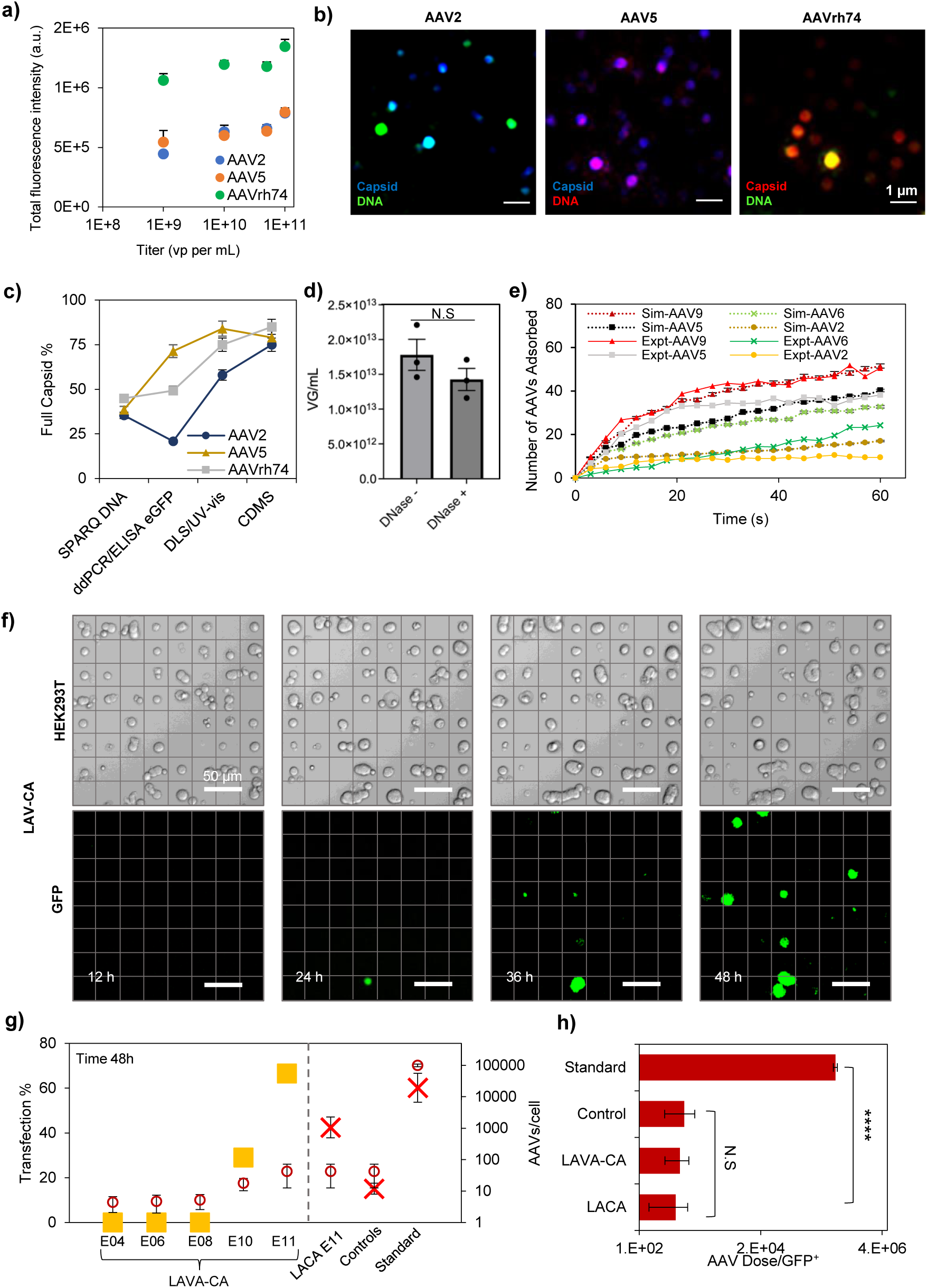
Serotype-dependent AAV capsid characterization and particle-number-limited single-cell transduction. Total fluorescence intensity measured for AAV2, AAV5, and AAVrh74 increases with input titer (10⁹-10¹¹ vgmL⁻¹), showing serotype-dependent differences in surface-associated signal at matched concentrations (n = 3, t-test). **(b)** Representative diffraction-limited images showing capsid (VP1-3, red) and genome-associated DNA (green) signals for AAV2, AAV5, and AAVrh74, illustrating serotype-specific particle distributions and capsid-genome co-localization at the single-particle level (scale bars, 1 µm). **(c)** Comparison of full capsid fraction across serotypes measured by SPARQ platform, ddPCR/ELISA-based infectivity, DLS/UV-Vis, and CDMS, showing overestimation of full particles by ensemble assays relative to diffraction-limited measurements (n = 3)**. (d)** The viral titers of the ssAAVrh74.CMV.eGFP.U6.milacZ were determined with (right) or without (left) DNase I treatment before lysis. Titers were presented as vector genome per milliliter (vgmL^-1^). The DNase I untreated samples had a viral titer of 1.78 ± 0.22 × 10^13^ vgmL^-1^, and the DNase I-treated samples had a viral titer of 1.43 ± 0.16 × 10^13^ vgmL^-1^. Following DNase I treatment, the ssAAVrh74 titer did not significantly decrease (*p*-value: 0.087, unpaired two-tailed t-test, n = 3). **(e)** Time-resolved adsorption kinetics of AAV2, AAV5, AAV6, and AAV9 in SPARQ, comparing experimental measurements (solid lines) with multi-physics simulations (dashed lines) (n = 9). **(f)** Time-lapse phase-contrast and eGFP^+^ fluorescence images of HEK293T cells immobilized by light-activated viral-cell adsorption (SPARQ) following localized AAV exposure, showing emergence of eGFP expression between 24-48 h post-exposure and confinement to patterned regions (scale bars, 50 µm). **(g)** Quantification of transduction efficiency at 48 h across decreasing AAV inputs on SPARQ (E4-E11; n = 3). **(h)** Bars represent the effective AAV dose required per eGFP^+^ cell across delivery modalities (n = 3; one-way ANOVA with Tukey’s HSD test; \*\*\*\**p* < 0.0001; N.S., not significant).

### ddPCR underestimates unpackaged AAV genomes

To determine whether ssAAVrh74.CMV.eGFP.U6.milacZ viral preparation contained unpackaged vector genomes resulting from degradation or the method used to produce the AAV. The preparation was treated with DNase I before lysis and compared its viral titer to that of an untreated control. DNase-treated samples yielded titers with a mean of 1.42×10^13^ vgmL^-1^ (S.D ± 7.57× 10^12^, n = 3), which were compared with untreated controls with a mean of 1.78×10^13^ vgmL^-1^ (S.D ± 4.06× 10^12^, n = 3). These results showed no statistically significant (p>0.05) decrease in viral titer in the DNase I-treated samples, indicating that the ddPCR method lacks sensitivity to detect unpackaged or degraded AAV vector genomes (**Fig**. **2d**).

### Serotype-dependent adsorption is governed by electrostatics

Real-time quantification of AAV adsorption with SPARQ revealed significant serotype-dependent binding kinetics that matched the multi-physics simulation trend. Fluorescently labelled fibronectin (Alexa Fluor™ 647) confirms the fidelity and spatial uniformity of surface patterning used for immobilization. Under identical initial bulk concentrations (1×10^8^ - 1×10^10^) and hydrodynamic conditions, AAV2 exhibited the slowest adsorption kinetics and the lowest steady-state occupancy (9.5 ± 4.04 AAVs/pattern, n = 4), followed by AAV6 (24.25 ± 2.98 AAVs/pattern, n = 4), whereas AAV5 (38.25 ± 7.41 AAVs/pattern, n = 4) and AAV9 (50.5 ± 14.24 AAVs/pattern, n = 4) displayed progressively faster accumulation and higher final AAV counts on the patterned surface. Experimental trajectories showed a biphasic kinetic profile, rapid initial adsorption during the first 10-15 secs, followed by a slower approach to saturation (40-60 secs), consistent with diffusion-limited transport transitioning to surface-interaction-limited binding. Multiphysics simulations accurately matched the experimentally observed adsorption curves across serotypes, with AAV2 (17 ± 3.36 AAVs/pattern, n = 4), followed by AAV6 (32.75 ± 2.36 AAVs/pattern, n = 4), AAV5 (40.25 ± 3.02 AAVs/pattern, n = 4) and AAV9 (51.5 ± 8.26 AAVs/pattern, n = 4), indicating that particle transport is governed by a balance of advective-diffusive transport, stochastic Brownian motion, and electrostatic interactions **(Fig. 2e)**. In the low-Reynolds-number regime (Re<1) relevant to these experiments, AAV motion is dominated by Brownian diffusion (D = 5.1×10^-12^ m^2^s^-1^) and viscous drag, with gravitational settling negligible (Bond number < 1). As AAVs approach the surface, electrostatic interactions between the negatively charged viral capsid and the positively charged SPARQ regions become the dominant force governing adsorption probability. The experimentally observed ordering of adsorption efficiency correlates strongly with capsid zeta potential. Viral particles with less negative surface charge experienced weaker electrostatic repulsion, higher terminal velocity near the surface, and a lower energetic barrier for surface capture, resulting in faster adsorption kinetics and higher steady-state occupancy. In contrast, near-neutral charged capsids like AAV2 (-3.5 mV) require greater diffusive residence time near the surface for electrostatic attraction to overcome thermal motion, leading to slower accumulation and reduced binding efficiency. These effects are explicitly captured in the simulations through charge-dependent electrostatic force terms superimposed on Brownian trajectories.

### SPARQ for threshold estimation of AAV dose for single-cell transduction

To directly link particle binding dynamics to functional gene expression, we immobilized individual HEK293T cells on UV micropatterned SPARQ devices and exposed them to spatially confined, known numbers of surface-bound AAV9s. Under these conditions, transgene expressions were monitored as a direct function of the number of viral particles physically available to each cell. eGFP expression emerged in a time-dependent manner, becoming detectable between 24 and 48 h post-exposure, and remained strictly localized to patterned regions, showing that transduction arose from controlled, surface-associated viral delivery rather than bulk diffusion **(Fig. 2f)**. Quantitative analysis of transduction efficiency across different AAV input concentrations revealed a threshold-governed response that depended on both viral availability and spatial distribution of cells. In the SPARQ-based light-activated viral-cell adsorption (SPARQ) condition, where both AAVs and cells were co-patterned, no detectable eGFP expression was observed at low input concentrations (1×10^4^- 1×10^8^; ≤ ∼5 ± 2.52 AAVs/pattern), indicating that a low number of AAVs was not efficient to drive productive transduction. At higher inputs, a significant transition emerged: at 1×10^8^ (∼18 ± 7.55 AAVs per pattern), transduction efficiency increased to ∼29% (S.D ± 3.31%), and further increased to ∼66% (S.D ± 1.66%) at 1×10^11^ (∼42 ± 9.69 AAVs per pattern). This sharp rise demonstrated the existence of a critical AAV-number threshold required for successful single-cell transduction. In contrast, when only cells were patterned with SPARQ-based light-activated cell adsorption, and 1×10^11^ AAVs were delivered in suspension (suspension), transduction efficiency was reduced (∼42% ± 4.69%) despite identical nominal AAV input per cell, indicating that spatial confinement and AAV availability at the cell-surface interface strongly influenced the functional delivery. Cells exposed to un-patterned AAVs under control conditions (cells and AAVs in suspension, ∼42 AAVs per cell) exhibited substantially lower transduction efficiency (∼15%), further highlighting that bulk-equivalent AAV numbers do not translate into equivalent functional dose at the single-cell level. Notably, the standard suspension-based infection protocol, employing 100,000 AAVs per cell, yielded ∼60% transduction efficiency, which is comparable to the SPARQ conditions using ∼42 AAVs per cell. When normalized to functional output, co-patterning reduced the effective AAV dose required per eGFP-positive (eGFP⁺) cell by more than three orders of magnitude relative to standard bulk delivery **(Fig. 2g)**.

Normalization of viral input to functional outcome enabled direct estimation of the effective number of AAVs required to achieve a single successful transduction event (eGFP⁺ cell) under each delivery condition. This analysis reframes the transduction efficiency as a probabilistic process, where the reported value represents the average AAV dose required to ensure at least one productive gene expression event with high confidence. In the co-patterning assay with AAVs and cells, the average number of AAVs required per eGFP⁺ event was found to be 590 (S.D ± 283 AAVs), despite spanning a wide range of input concentrations (1×10^4^- 1×10^11^). Notably, this value was lower than that observed when only cells were patterned, and AAVs were delivered in suspension (493, S.D ± 339 AAVs per eGFP⁺ event), indicating that spatial confinement of viral particles at the cell interface substantially enhances the probability of productive infection even in the absence of viral patterning. In contrast, control conditions in which both cells and AAVs were delivered in suspension at a nominal dose of ∼42 (S.D ± 9.69) AAVs per cell required 716 (S.D ± 412 AAVs) AAVs to achieve a single eGFP⁺ event, reflecting inefficient AAV-cell encounters under bulk delivery. Importantly, the standard suspension-based infection protocol employing 100,000 AAVs per cell required approximately 52,000 (S.D. ± 42,458 AAVs) AAVs to reliably generate one eGFP⁺ cell **(Fig. 2h)**, showing a >100-fold increase in required viral input relative to the SPARQ assay.

## Discussion

We introduce SPARQ to define the functional AAVx dose by directly quantifying its availability, integrity, and transduction outcome at the single-particle and single-cell levels. Existing AAVx titration and characterization methods, including qPCR/ddPCR, capsid ELISA, AUC, SEC-MALS, CDMS, and bulk infectivity assays, measure ensemble-averaged properties and therefore mask particle-level heterogeneity in capsid integrity, genome retention, and biological competence^8,10,20,22,37^. Even advanced bulk approaches that resolve empty and full capsids remain decoupled from functional outcomes at the level of individual cells^15,24^. By contrast, SPARQ integrates viral adsorption, diffraction-limited multi-plane imaging, and controlled particle mapping to directly count how many physical AAVx particles engage a single cell and how many of those interactions culminate in productive gene expression. From a physics perspective, this reframes AAVx transduction as a stochastic, force-governed process in which diffusion, electrostatic interactions, and surface capture probabilities define local viral density, while gene expression emerges only when probabilistic barriers are exceeded^33,38^. Thus, SPARQ serves as a quantitative particle-cell interaction assay rather than a proxy for bulk dosing.

A major advantage of SPARQ is the experimental determination of a minimum AAVx number threshold for effective single-cell transduction, providing a direct and operational definition of effective AAVx MOI. Our data demonstrates that productive gene expression does not arise from isolated viral vector encounters but instead requires a critical local AAVx density, in the order of tens of particles per cell, above which transduction probability increases sharply and then saturates. This observation supports the hypothesis that AAVx density, rather than nominal bulk MOI, is the dominant determinant of transduction efficiency in adherent cell systems^11,39,40^. Conventional bulk assays are intrinsically limited in this regard, as stochastic and spatially uneven AAVx distribution across cells leads to underestimation of transduction efficiency and overestimation of neutralization by antibodies or serum factors^41–43^. By ensuring that each patterned cell is exposed to a known and quantified number of AAVx particles, SPARQ measures functional doses at single-cell resolution. This spatial confinement ensures that viral particles remain localized at the cell-surface interface, increasing the probability of virus-cell encounters and reducing losses associated with diffusion, dilution, or nonspecific adsorption. In contrast, suspension-based delivery relies on stochastic diffusion-mediated encounters between viruses and cells, where most viral particles fail to reach the cell membrane within a practical timeframe. These observations highlight how spatially controlled viral presentation fundamentally alters the efficiency of gene delivery compared with conventional bulk infection protocols. Importantly, prior studies addressing the number of AAVx genomes expressed per cell relied on indirect inference from delayed reporter co-expression and probabilistic modeling rather than direct quantification of incoming particles^44^. SPARQ overcomes this limitation by directly counting AAVx particles upstream of expression, thereby providing strong co-relation between physical particle number and functional outcome. This technology also offers a physical rationale for delivery strategies that increase local AAVx density, such as encapsulation of multiple AAVx within extracellular vesicles, where a single EV carrying ∼20-40 AAVx could function analogously to a SPARQ pattern that reliably produces a transduction event at substantially reduced systemic dose^3,45,46^.

Beyond defining functional dose, SPARQ has important implications for clinical translation, particularly in the context of AAVx neutralization and patient eligibility for gene therapy. Current neutralizing antibody (NAb) assays rely on bulk inhibition of transduction and are confounded by assay sensitivity, uneven viral distribution, and the presence of non-neutralizing but binding anti-AAVx antibodies, leading to frequent overestimation of neutralization potency^47–49^. As a result, many patients are excluded from gene therapy despite retaining partial or full transduction capacity in vivo^41,50,51^. By quantifying transduction probability as a function of a precisely defined particle number at the single-cell level, SPARQ estimates a more accurate assessment of true functional neutralization. While the current implementation is optimized for in vitro patterned substrates and high-resolution imaging, these limitations are addressable through automation, higher-throughput patterning, and integration with microfluidic and organoid platforms. Future extensions of this approach could enable comparative evaluation of engineered capsids, EV encapsulated AAVx, and antibody-virus complexes under controlled particle-density conditions. More broadly, this work establishes a generalizable technique for defining viral dose, integrity, and function as a coupled physical process that may ultimately improve the safety, efficiency, and accessibility of AAVx gene therapies.

## Conclusion

SPARQ is a new high-throughput AAVx quality assessment tool for defining critical functional AAVx dose estimation. This study redefines characterization of AAVx by demonstrating that transduction is governed by the local availability of a discrete number of viral particles at the single-cell level, rather than by bulk vector concentration. By directly quantifying particle-cell interactions and linking them to gene expression, we show that productive transduction emerges only above a critical particle-number threshold that is orders of magnitude lower than doses used in conventional suspension-based protocols. Clinically, this technology enables rational dose minimization, more accurate interpretation of neutralization assays, and improved patient stratification. By grounding AAV delivery in a quantitative, physics-informed model of viral engagement, this work provides a path toward safer, more efficient, and more precise gene therapy.

## Acknowledgments

Electron microscopy was performed at the Center for Electron Microscopy and Analysis (CEMAS) at The Ohio State University. Computations were performed with the help of the Ohio Supercomputer Center, Columbus, Ohio.

## Funding

This work was supported by the U.S. National Institutes of Health (NIH) grants R21TR005565 (E.R. & N.Y.S.), and institutional funds from Nationwide Children’s Hospital (S.Q.H. & N.Y.S.). Additional support for E.R. was provided by the College of Engineering (COE) Strategic Research Initiative Grant, the William G. Lowrie Department of Chemical and Biomolecular Engineering and the James Comprehensive Cancer Center at The Ohio State University.

## Author contributions

E.R. conceived the idea. E.R., N.Y.S., C.K.N, and K.T.N. developed experimental plans and designed the study. C.K.N and K.T.N performed experiments. C.K.N, E.R., and N.Y.S. drafted the manuscript with input from all authors. K.J.K. designed probes and controls used in the study. S.Q.H. provided AAV vectors. C.K.N and K.T.N prepared tables and figures. B.B. and J.N. performed AAV bulk analytical assays. X.H prepared cells for transduction and engineered micropatterns for SPARQ. X.Y.R. adapted a MATLAB algorithm for single-particle and colocalization detection, engineered micropatterns and supervised transduction study. J.D.R performed purification for antibody-AAV labelling studies. E.R supervised the whole study and edited the manuscript. All authors provided critical feedback and helped shape the research, analysis, and manuscript.

## Competing interests

E.R., K.T.N., C.K.N., and N.Y.S. have filed a provisional patent application for the SPARQ platform.

## Data and materials availability

All data needed to evaluate the conclusions in the paper are presented. Additional data related to this paper may be requested from the authors.

## Notes

### Competing Interest Statement

Authors: Eduardo Reategui., Kim Truc Nguyen., Chiranth K Nagaraj., and Nizar Y Saad. have filed a provisional patent application for the SPARQ platform.

